# Sensorimotor Impairment in Ageing and Neurocognitive Disorders: Beat Synchronisation and Adaptation to Tempo Changes

**DOI:** 10.1101/2023.12.27.573413

**Authors:** Andres von Schnehen, Lise Hobeika, Marion Houot, Arnaud Recher, François Puisieux, Dominique Huvent-Grelle, Séverine Samson

## Abstract

**Background:** Understanding the nature and extent of sensorimotor decline in ageing individuals and those with neurocognitive disorders NCD, such as Alzheimer’s disease, is essential for designing effective music-based interventions.

**Objective:** Our understanding of rhythmic functions remains incomplete, particularly in how ageing and NCD affect sensorimotor synchronisation and adaptation to tempo changes. This study aims to fill this knowledge gap.

**Methods:** Patients from a memory clinic participated in a tapping task, synchronising with metronomic and musical sequences, some of which contained sudden tempo changes. After exclusions, 51 patients were included in the final analysis.

**Results:** Participants’ mini-mental state examination scores were associated with tapping consistency. Additionally, age negatively influenced consistency when synchronising with a musical beat, whereas consistency remained stable across age when tapping with a metronome.

**Conclusions:** The ability to extract a beat from a musical signal diminishes with age, whereas the capacity to maintain a beat remains relatively constant. However, both processes may decline at moderate or severe stages of NCD. Moreover, the results indicate that the initial decline of attention and working memory with age may impact perception and synchronisation to a musical beat, whereas progressive NCD-related cognitive decline results in more widespread sensorimotor decline, affecting tapping irrespective of audio type. These findings underline the importance of customising rhythm-based interventions to the needs of older adults and individuals with NCD, taking into consideration their cognitive as well as their rhythmic aptitudes.

This study was registered at clinicaltrials.gov (NCT04146688).

## INTRODUCTION

### Neurocognitive disorders and music-based interventions

Neurocognitive disorders (NCD) are acquired disorders marked by a progressive decline in cognitive functioning, particularly with regards to memory, but also including domains like attention, language, learning, and social cognition, challenging the patients’ capacity to live autonomously [1]. Before the fifth edition of the Diagnostic and Statistical Manual of Mental Disorders (DSM-5), major NCD was referred to as dementia, but we will use the term NCD in this article. Different forms exist, such as Alzheimer’s disease (AD), vascular NCD, NCD with Lewy bodies, and others. In the same vein, minor NCD is a DSM-5 diagnosis corresponding to a milder or prodromal form of the disease, which generally does not impede autonomy, and which was referred to as mild cognitive impairment before. In the absence of a cure for NCD, there is promise in improving the quality of life of those affected by enhancing various aspects of their well-being through non-pharmacological interventions. Among these interventions, music-based approaches have shown considerable potential in this regard. It has been suggested [2–4] that they may be particularly effective if they stimulate sensorimotor synchronisation (SMS), defined as the temporal coordination of rhythmic movement with an external rhythm [5]. This may be related to temporal expectations elicited by a musical beat, which may stimulate the reward network and induce pleasure [6]. Besides directly eliciting reward and pleasure, stimulating rhythmic abilities may have positive effects on the way people with NCD interact with and adapt to their environment. Improving temporal prediction abilities might help people synchronise and interact with others [7], improve communication, and reduce isolation.

### Sensorimotor synchronisation

In SMS research, individuals coordinate their movements with an auditory sequence, typically involving a simple metronome or music. Synchronisation performance is typically assessed in terms of consistency and asynchrony. Consistency refers to the degree of variability in the time differences between taps and beats, whereas asynchrony refers to whether participants tapped before (negative asynchrony) or after (positive asynchrony) the pacing event [8]. Paced tapping to a metronome and paced tapping to music, while ostensibly the same task, may in fact engage different mechanisms. In a metronomic sequence, the beat is indicated as simple regular tones, whereas in music, the beat is embedded within a complex auditory pattern. In this context, it is useful to think of beat perception as being comprised of two subprocesses; beat induction (beat finding), where an underlying beat is inferred even when auditory events are unequally spaced [9,10]; and beat maintenance (beat continuation), which is a more implicit and mechanical process that implies continual, sustained measurement of predictable intervals and is less dependent on attention [10,11]. Likely, both processes are employed simultaneously, the relative dependence of each depending on the saliency of the beat. Tapping to a metronome, then, might employ primarily beat maintenance processes, whereas tapping to music may be more dependent on beat induction. As a result, tapping to music is often associated with higher difficulty, expressed in lower tapping consistency [12–14], but not all studies have confirmed this [8,15]. Indeed, the difficulty of performing synchronous movement to music presumably depends primarily on the clarity of its beat. In terms of asynchrony, people tend to tap ahead of the beat (referred to as mean negative asynchrony) when synchronising with a metronome but not with music [8,12–14,16]. The mechanisms underlying this phenomenon are still not fully understood.

A key process in SMS is error correction. While error correction is an ever-present mechanism without which one would gradually become out of sync [17], it can directly be tested by introducing tempo changes. Adapting to tempo changes requires attention, awareness and some memory for at least the preceding events [18], and is likely related to cognitive flexibility, the ability to shift between mental sets and strategies [19].

### How do age and neurocognitive disorders impact sensorimotor synchronisation?

#### Ageing

The ability to perceive a beat and synchronise to it emerges early in life, remains relatively stable in adulthood [20,21], and may be preserved in old age, at least when synchronising with an evenly spaced beat at a comfortable tempo [22–25]. Nonetheless, certain studies have indicated a decline of sensorimotor abilities associated with age, which seems to appear above the age of 75 [20,21,26] (however, see [27] for a study demonstrating a reduction in tapping performance even in relatively young older adults). However, there remains a lack of research studying SMS in the latest decades of life. Importantly, another study [28] did not find age-related differences in simple SMS, but older participants’ performance was diminished when participants had to simultaneously perform a cognitively challenging task while tapping to a metronome. This suggests that older people may employ more attention and working memory resources when tapping at a comfortable tempo.

The aforementioned studies examined the effect of age on SMS using metronomes. To our knowledge, the influence of old age on SMS with *music* has not yet been tested. However, considering the typical decline in attentional capacities associated with ageing [29,30], it is reasonable to speculate that beat induction may be more vulnerable to age-related decline than beat maintenance, which would result in age-related declines in SMS performance particularly with music.

#### Neurocognitive disorders

Several studies have indicated that SMS abilities tend to be relatively preserved in individuals with NCD when instructed to tap along with a metronome set to a comfortable tempo [31]. However, differences were observed when participants had to continue tapping after an external sequence had ended [22,32] or when the target rate deviated substantially from their comfortable tempo [22,33,34], manipulations likely to engage working memory and attention. However, a recent study by Hobeika et al. [35] revealed a decline in performance among individuals with major NCD in tapping even at a comfortable rate with a metronome and with music, and a negative relationship between consistency during metronome tapping and participants’ score on the mini-mental state examination (MMSE) [36], a brief screening tool for assessing NCD. These different results may be explained by the severity of NCD in Hobeika et al.’s study (their NCD group had a mean MMSE of 15.5, which was lower than in the other studies). However, definite conclusions cannot be drawn from Hobeika et al.’s study because they pooled together participants who were tested under different conditions, which emphasises the need for the current study, investigating the impact of NCD severity under homogeneous testing conditions.

It may be that in healthy ageing people, or mild forms of NCD, tapping with a metronome may be a largely automatic process, and only when tapping with music, or in cognitively challenging paradigms are attention and working memory processes recruited, leading to a decline in performance. However, for people at more severe stages of NCD, attention and working memory are needed even to synchronise at a comfortable pace with a metronome. Indeed, some research has demonstrated increased use of non-motor regions during simple motor tasks in NCD, at least in AD [37,38], indicating that with increasing severity of the disease, individuals employ other domain-general brain networks during SMS, reflecting a shift to more effortful and less automatic processing of rhythm, and potentially lower performance due to increasing competition for limited cognitive resources. It is possible that individuals in moderate to severe stages of NCD may require more attention and working memory even for tapping with simple metronomic sequences, and may be more impaired in these cognitive abilities, resulting in a general sensorimotor impairment manifested by reduced consistency when tapping to any regular stimulus, be it a metronome or music.

Similar to synchronisation-continuation tapping and tapping at tempi far from one’s comfortable rate, tapping with a sequence containing tempo changes may particularly involve attention and working memory [18]. A particular impairment in SMS with tempo changes has already been demonstrated in other clinical populations, namely, people with traumatic brain injury [39], autism spectrum disorder [40], basal ganglia pathology [41], and cerebellar lesions [42], and has been explained in terms of attention-dependent temporal processing. Since attention is greatly impaired in people with NCD [43], the ability of people with NCD to adapt their tempo when encountering tempo changes may be compromised, due to an imprecise representation of temporal structure and inefficient allocation of attention over time [41]. Finally, a decline in cognitive flexibility found in people with NCD [44,45] may present another contributing factor to their potential disadvantage in adaptation to tempo changes. To our knowledge, there does not exist any research examining the effect of NCD on tempo adaptation in SMS. It is important to investigate this aspect, as adaptation to tempo changes can serve as a model for understanding how individuals interact with a dynamically changing environment in general.

### The current study

The aim of the current study was to test the effects of age and NCD severity on SMS to metronomes and to music with and without tempo changes, in a group of patients at a French memory clinic, most of whom had major or minor NCD of diverse aetiologies. In this context, we examined SMS skills, with particular emphasis on the impact of tempo changes by introducing sudden accelerations and decelerations every 15 seconds in half of the trials, and computing consistency and asynchrony. The difficulty in this task should come only from the changes in tempo, rather than presenting participants with inherently difficult tempi for synchronisation. To achieve this, we selected base tempi that closely aligned with the typical spontaneous motor tempo reported in the literature for older adults [21,25] and we confirmed this by assessing individuals’ spontaneous tempo. In traditional SMS paradigms, participants typically tap their finger or hand to an auditory regular beat. However, when applying such paradigms to individuals with NCD, particularly in advanced stages, special consideration is required to avoid stressful, unpleasant or artificial laboratory situations, as they might find it difficult to cope with such conditions, and they may experience difficulties in retaining and following instructions, especially in longer experiments. For these reasons, tasks with multimodal stimuli, creating a quasi-social situation, may be conducive [46,47]. Additionally, people, including older adults with NCD, might actually perform better when synchronising with a video than with another person [14,48]. We recently developed and validated an experimental setup tailored to elderly individuals [14,48,49] and continue its use to present stimuli bimodally (audio plus video). Using this experimental design and assessing a group of older adults exhibiting a range of ages and varying levels of cognitive impairment created an optimal setting for examining the distinct impacts of both age and NCD on sensorimotor synchronisation.

#### Hypotheses

Firstly, we expect a global impairment in SMS with increasing NCD severity. Specifically, we expect that MMSE score will have a negative impact on tapping consistency. Additionally, we hypothesise that consistency will be lower in trials with a shifting tempo compared to those with a stable tempo. More importantly, we predict an interaction between the presence or absence of tempo changes and MMSE score, such that the reduction in consistency in the shifting condition will be more pronounced in individuals with a lower MMSE score, probably due to declines in attention, working memory, and cognitive flexibility. Consistent with previous research, we expect lower consistency when individuals synchronise their movements with music compared to a metronome [12–14]. Furthermore, due to increased reliance on beat induction with music and decreased attentional capacities with ageing, we expect consistency to decrease with age when individuals tap with music, but to a lesser extent (or not at all) with a metronome. Finally, we hypothesise that asynchrony will be lower (more negative) in the metronome conditions compared to music [5,8,12–14,50].

## MATERIALS AND METHODS

### Participants

A total of 61 patients were recruited at the geriatric day hospital *Les Bateliers* (Lille University Hospital, France), during a scheduled consultation related to memory problems or falls. Inclusion criteria included age between 60 and 99, right-handedness and native or near-native fluency in French. Patients were ineligible for participation if they had Parkinson’s disease, other motor disorders or paralysis, or uncorrected hearing or vision problems. Patients’ data were excluded from analysis if they did not finish the experiment. Their diagnosis of major NCD, minor NCD, or absence of NCD was made by a geriatrician and based on DSM-5 criteria [1]. However, in this study, we assessed cognitive impairment as a continuous variable using the MMSE. After ten exclusions (seven who withdrew from the study during the experiment, one due to technical problems, one who tapped in a seemingly random fashion as indicated by Rayleigh’s test [51], and one whose MMSE score of 14 was an outlier; 3 SDs below the mean), 51 patients were included in this study. The data were collected between November 2021 and July 2022. The study was approved by the local Ethics Committee (Comité de Protection des Personnes, Sud-Est VI, France; No. 2017-A03543-50) and by the Commission Nationale de l’Informatique et des Libertés, registered at clinicaltrials.gov (NCT04146688). All patients provided written informed consent for their participation in accordance with the Declaration of Helsinki.

### Materials

#### Experimental apparatus

The experimental set-up consisted of a chair for the patient with a tapping tablet attached to the right armrest [46,47]. A life-sized screen (158 x 92 cm) and a pair of loudspeakers were placed in front of the patient at a distance of 230 cm. A video of a musician tapping to the simultaneously presented auditory sequence was projected onto the screen during the task in front of the patient. Each patient was tested individually and was separated from the experimenter by a curtain to avoid distraction. Stimuli were presented and responses collected using a programme written in MAX/MSP (https://cycling74.com).

#### Stimuli

Stimuli were 75 seconds long and consisted of either a metronome or a musical sequence and a video recording of the musician tapping to the beat of the auditory sequence. Both types of audio were preceded by 4 beats to provide the tempo.

Metronome trials consisted of regular beats. For the music trials, a custom-made rendition of an excerpt of the French popular song “Non, je ne regrette rien” by Édith Piaf was used. This particular song was chosen because it was likely well-known to our age group and its original tempo is close to older adults’ spontaneous motor tempo [14,25]. A MIDI version of the song (without lyrics), available in an online music repository (www.midis101.com) was selected and cropped to a length of 75 s. We opted for a MIDI version of the song in order to have completely isochronous timing and the possibility to manipulate its tempo.

The musical and metronomic sequences were manipulated to conform to one of four temporal patterns (Figure 1): A stable IOI of 674 ms (A), a stable IOI of 741 ms (B), or a sequence in which the tempo shifted every 15 seconds between the two, starting either at 674 ms (C) or starting at 741 ms (D). Ableton Live was used to render the musical stimuli from the MIDI versions using their in-house instruments, to create the metronomic stimuli and to perform the tempo manipulations. The visual part of the audio-visual stimuli was created beforehand by filming the musician who sat in the position of the participants, listened to the musical stimuli and tapped along. An analysis of the musician’s tapping consistency and asynchrony during the recording of these videos indicated very good performance and minimal error (see Table S1).

**Figure 1.**
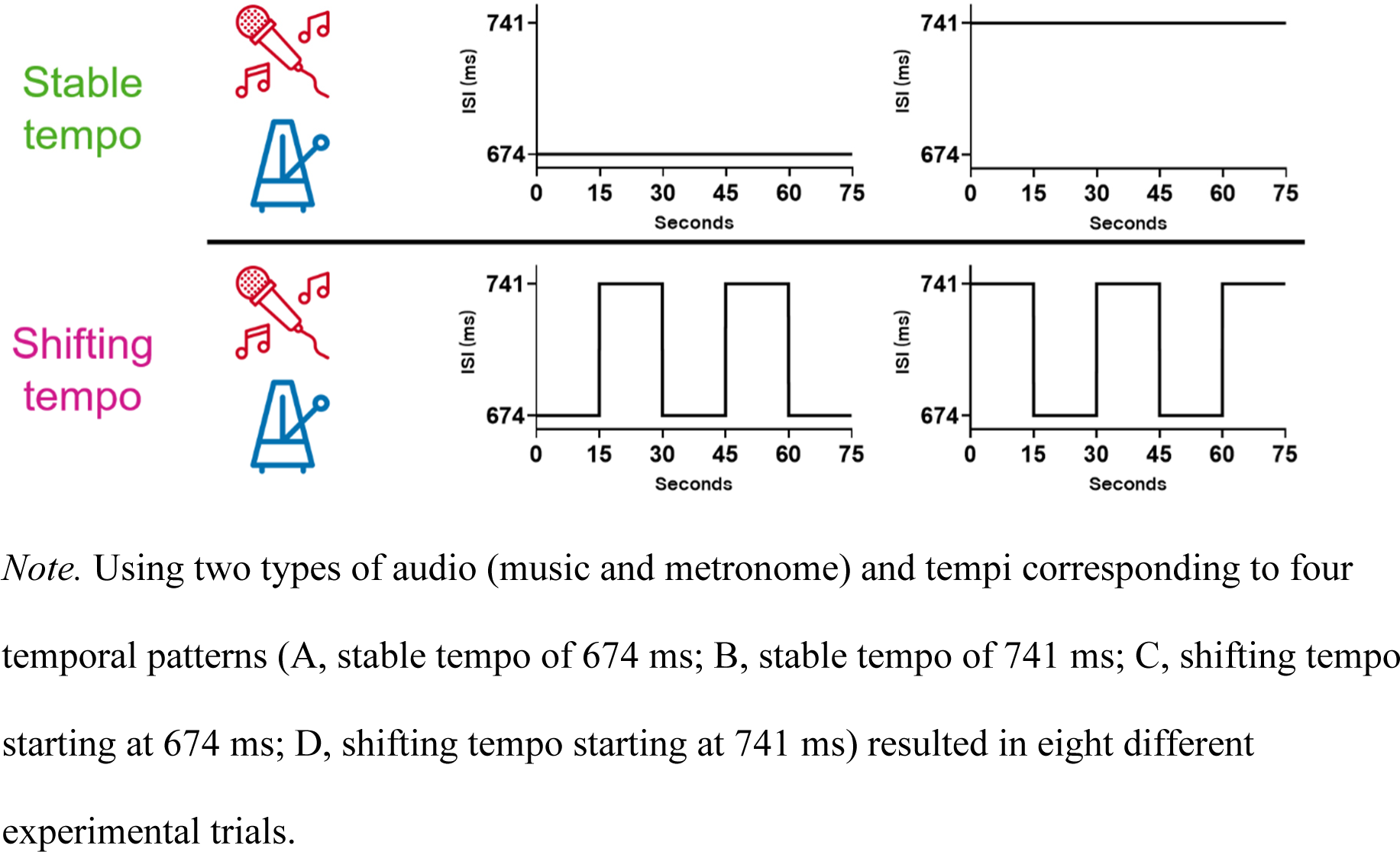
Types of audio and temporal structures used in the experimental trials

### Procedure

The experiment started by orally administering a musical expertise questionnaire, which inquired about participants’ musical training, listening habits, and engagement with music. Then, short forms of the Geriatric Depression Scale [52] and the Geriatric Anxiety Inventory [53] were orally administered. Next, each patient performed a brief spontaneous motor task by tapping as regularly as possible for 31 taps (30 inter-tap intervals; ITIs), at their preferred, comfortable tempo.

Afterwards, each patient underwent the paced tapping task, in which they were exposed to bimodal stimuli (described in the preceding section) and tapped along with every beat, just like they watched the musician do in the video. A practice trial was followed by eight experimental trials, counterbalanced across participants, in a randomized order. The participant was not informed that tempo changes might occur. The patient was given the possibility to take a break after half of the experimental trials.

### Data analysis

#### Calculation of SMS variables

For the 30 intervals produced during the spontaneous motor task, we calculated mean ITI and CV (standard deviation divided by mean) of ITI. In the paced tapping task, as we mentioned above, the tempo either remained stable or shifted every 15 seconds. This allowed us to analyse responses per 15-second segment to explore the impact of three within-subjects variables, each with two levels: audio (music/metronome), tempo (fast/slow), and tempo stability (stable/shifting; Figure S1A).

Consistency and asynchrony were computed using circular statistics [54] with the CircStat toolbox [55] in MATLAB [56]. We opted for circular analysis of synchronisation data as this allowed for a robust analysis even in the case of missing or superfluous taps, as asynchronies and their variability can be computed without necessarily attributing each response event to a particular beat [57]. In a given trial, ms in an inter-onset interval (IOI) are converted into degrees on a circular scale going from -180° to +180°. The beat’s onset is at 0°, the time a participant would be expected to tap. An angle of 180° would indicate a participant tapping in antiphase. Vectors were averaged to obtain a mean resultant vector R^-⃗^ [54,55] allowing for the calculation of synchronisation consistency and asynchrony. Consistency is represented by the length of the vector R^-⃗^ and ranges from 0 to 1, where 1 corresponds to perfect consistency (all taps occurred with the same delay to the beat) and 0 describes a situation where taps were randomly distributed between the beats. Asynchrony reflects the angular deviation (θ) of vector R^-⃗^ from 0, which is then transformed back into ms (Figure 2). Consistency and asynchrony were only analysed for the segments 2 through 5, as performance in the first segment was not pertinent to us since no tempo change would have occurred in this segment, even in the shifting condition (see Figure S1B).

**Figure 2.**
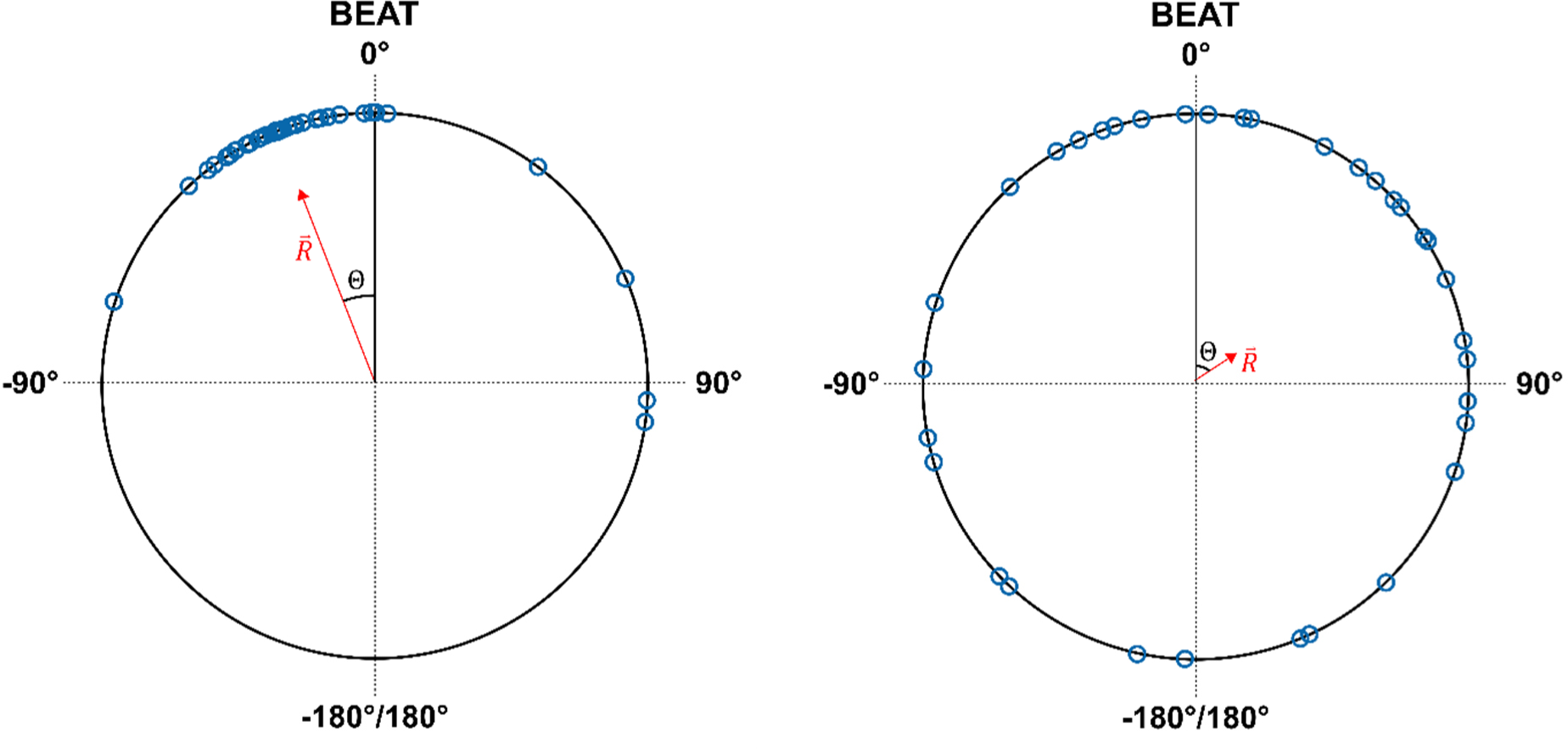
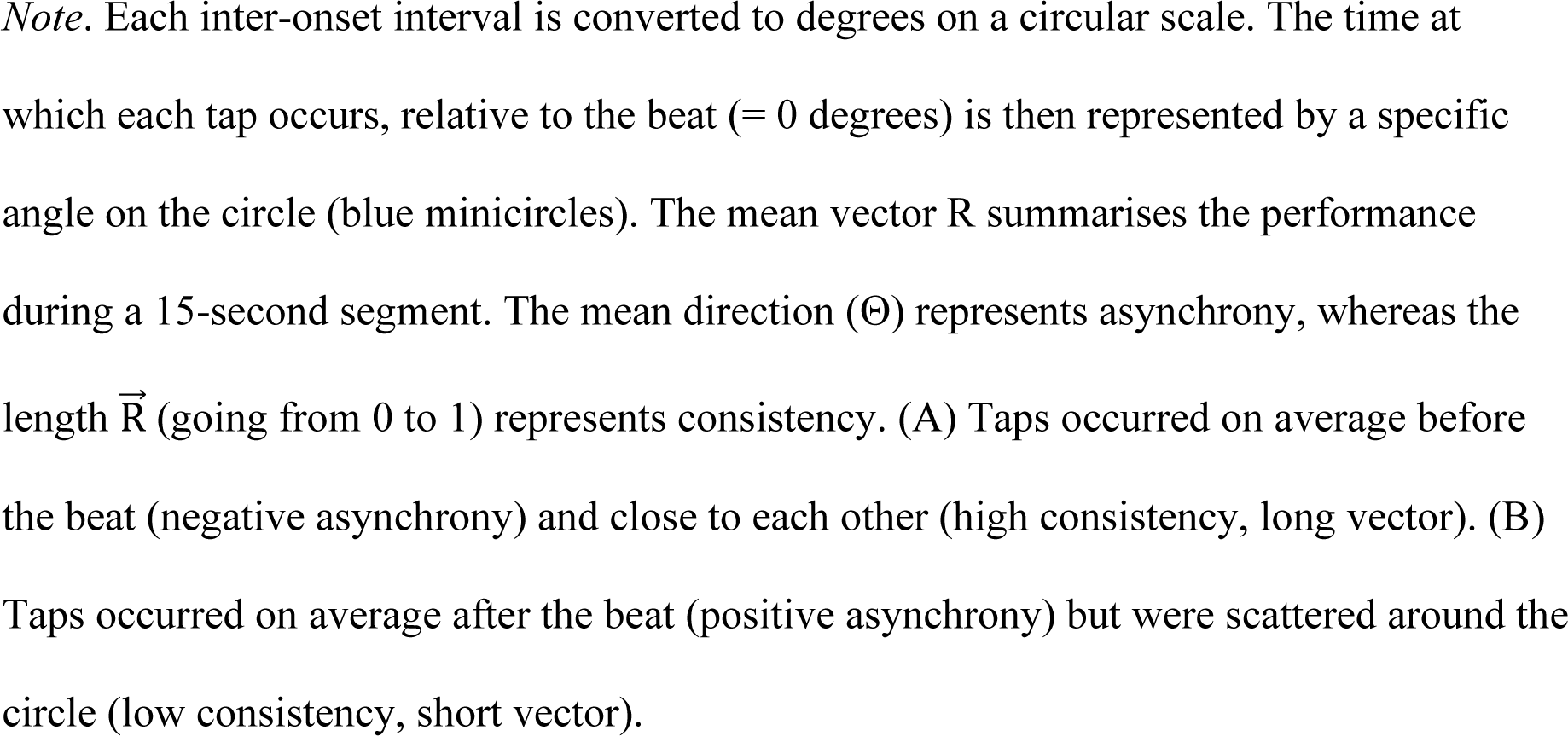
Two examples of circular synchronisation analysis in a given trial

#### Statistical analyses

All statistical analyses were performed in R 4.2.2 using RStudio [58,59]. We analysed SMS consistency and asynchrony by conducting mixed-effects models. In both cases, fixed effects included the four within-subject factors (audio, tempo, tempo stability, and segment) as well as the between-subjects variables age and MMSE. We added to both models the four-way interaction between audio, tempo, tempo stability, and segment, the four-way interaction between audio, tempo, tempo stability, and age, the four-way interaction between audio, tempo, tempo stability, and MMSE, as well as all lower-order interactions and main effects. Furthermore, we controlled for the effects of gender, years of education, musical expertise, and condition order by entering them as additional fixed effects in the model. Finally, participant was entered as a random effect. In the analysis with consistency as a dependent variable, a generalised linear mixed model with a beta distribution and a logit link was performed using the glmmTMB package [60] in R. In the analysis with asynchrony as a dependent variable, we first filtered out segments with insufficient taps (i.e., where the percentage of taps relative to the number of beats was more than 2 standard deviations below the mean). Then, we transformed the variable asynchrony by taking the cubic root of its absolute value and multiplying it with its original sign. This was done to fulfil the assumption of normality of residuals, as asynchrony was right-skewed. Then, we performed a linear mixed-effects model analysis using the lme4 package [61]. Type II Wald chi-square tests were used to test the main effects and interactions. We present effect size by computing f², which is considered an appropriate metric of effect size in mixed-effects regression models [62]. Only the significant highest-order interactions are presented and discussed even if they contain lower-order significant effects; however, the complete results of the two analyses are detailed in Tables S3 and S4. None of the significant effects including segment will be presented in the results section to focus on the effects of interest.

## RESULTS

### Participants

Demographic data, including age, gender, and education, and clinical data, encompassing diagnosis, MMSE, activities of daily living [63], instrumental activities of daily living [64], Geriatric Depression Scale, and Geriatric Anxiety Inventory, can be found in Table 1. Of the 21 participants diagnosed with major NCD, nine were diagnosed with AD, two with vascular NCD, 11 with NCD of mixed aetiology, and one with NCD of an unknown origin. A distribution of MMSE scores is shown in Figure S2. Demographic and clinical data of the seven participants who withdrew from participation during the study can be found in Table S2. The participants who did not finish the study were on average older (U = 66.0, *p* = .007, Mann-Whitney U test) and had a lower MMSE score (U = 265.5, *p* = .038, Mann-Whitney U test) than those who did, whereas the two groups did not differ in terms of gender, diagnosis, education, musical expertise, ADL, IADL, depression, or anxiety (all *p* > .05). Moreover, all participants who finished the study prematurely were diagnosed with major NCD. To ensure that the tempo of the experimental stimuli (674 ms and 741 ms) was within a comfortable range, participants’ mean spontaneous motor tempo was computed, which was 715 ms (SD = 468 ms).

**Table 1.** Demographic and clinical information of patients

|  | N | Median [first quartile, third quartile] or frequencies (%) | Range |
| --- | --- | --- | --- |
| Age | 51 | 82 [76, 86] | 61-92 |
| Gender (women) | 51 | 35 (69%) |  |
| Years of education | 51 | 12 [7, 14] | 3-18 |
| Musical expertise (out of 28) | 51 | 3 [2,4] | 0-12 |
| Diagnosis | 48 |  |  |
| Major NCD |  |  |  |
| Minor NCD |  |  |  |
| No NCD |  |  |  |
| MMSE (out of 30) | 51 | 25 [23, 28] | 19-30 |
| ADL (out of 6) | 50 | 5.5 [4.5, 6] | 2-6 |
| IADL (out of 4) | 50 | 2 [1, 3] | 0-4 |
| GDS (out of 15) | 50 | 5 [3, 8] | 1-12 |
| GAI (out of 5) | 50 | 2 [1, 4] | 0-5 |

NCD = neurocognitive disorder; MMSE = Mini-Mental State Examination; ADL = activities of daily living; IADL = instrumental activities of daily living; GDS = Geriatric Depression Scale; GAI = Geriatric Anxiety Inventory

### Consistency

The results of the generalised linear mixed model are presented in Table S3. A main effect of MMSE (Wald χ² = 4.06, *p* = .044, f² = 0.03) suggests that more cognitively impaired people (i.e., with a lower MMSE score) tapped with a lower level of consistency (Figure 3A). Furthermore, there was a significant interaction of audio and age (Wald χ² = 7.06, *p* = .008, f² < 0.01; Figure 3B). The slope of music was negative and significantly different from zero (*p* = .046), whereas the slope of metronome was not (p = .433). In other words, consistency decreased with age, but only in the music conditions.

**Figure 3.**
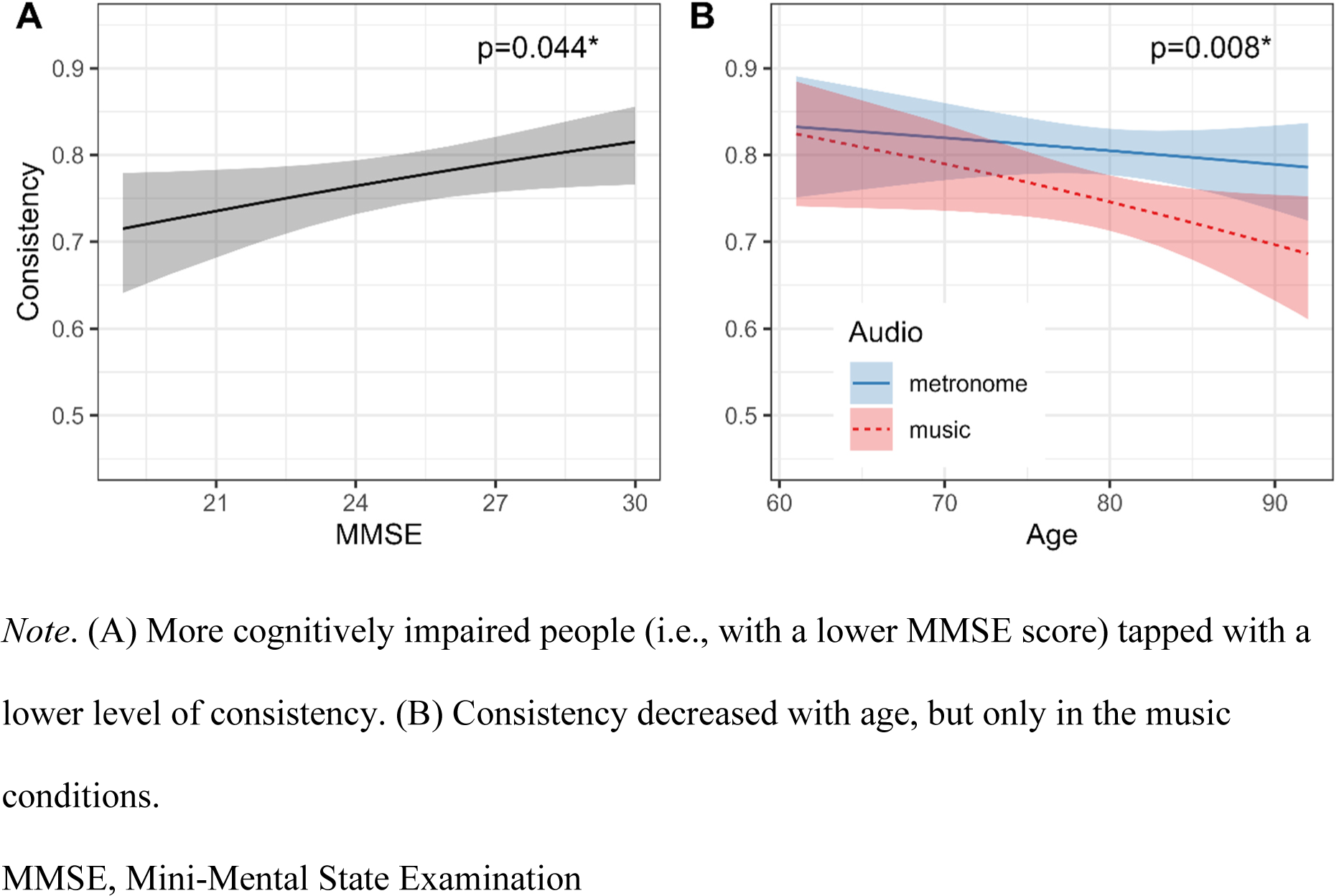
Effects of MMSE, and of the interaction of audio and age on consistency

Additionally, there was a three-way interaction effect of audio, tempo, and tempo stability (Wald χ² = 0.02, *p =* < 0.001; Figure 4A). We conducted post hoc tests for the 12 relevant pairwise comparisons: Conditions were compared in which two of the variables remained constant while the third varied (e.g., music/fast/stable versus music/fast/shifting). Statistical significance was adjusted for multiple comparisons using the Benjamini-Hochberg method. Across conditions, consistency was higher in metronome compared to music trials, and in trials with a stable tempo compared to those with a shifting tempo. Regarding tempo, consistency did not differ between segments with a fast versus slow tempo, with one notable exception: In metronome trials with a shifting tempo, consistency was higher when the tempo was slow (i.e., following a deceleration) compared to when it was fast (i.e., following an acceleration; mean difference estimate ± standard error: 0.03 ± 0.03). Finally, there were a two-way interaction effect of tempo stability and age, and a three-way interaction between audio, tempo, and MMSE. These effects were the two smallest effects (f² < .002) and it is unlikely they have practical significance. We therefore report these effects only in the Supplementary Information (Figures S3 and S4).

**Figure 4.**
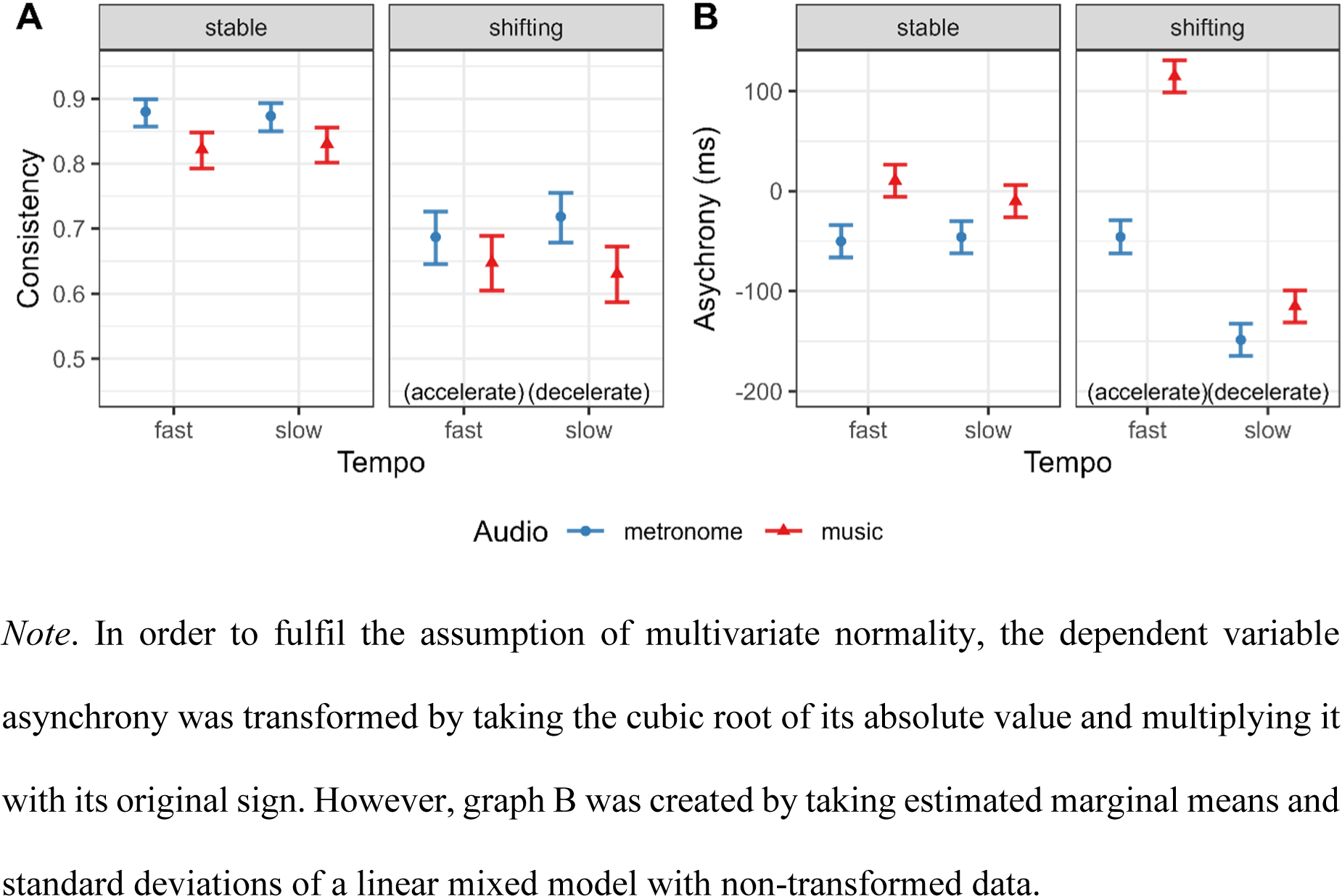
Effects of audio, tempo, and tempo stability on consistency (p = .107; A) and asynchrony (p < .001; B)

### Asynchrony

The results of the linear mixed model are presented in Table S4. There was a significant three-way interaction between audio, tempo, and tempo stability (Wald χ² = 15.15, *p* < .001, f² = 0.02; Figure 4B). As in the analysis of consistency, pairwise comparisons were conducted and statistical significance adjusted using the Benjamini-Hochberg method. All of the 12 pairwise comparisons yielded statistically significant results (p < .05), indicating significant differences among the means. When the tempo was stable, participants anticipated the stimuli more when tapping with a metronome than with a musical beat. When tapping to a sequence with a shifting tempo, we appear to see the interaction of two phenomena. On the one hand, people’s taps generally tended to occur somewhat later (higher or positive asynchrony) after an acceleration and somewhat before (lower or negative asynchrony) after a deceleration, which is to be expected: After a perturbation, participants’ taps may still correspond to the pre-perturbation tempo before they adapt to the changed tempo. On the other hand, just like when tapping with a stable tempo, people’s responses when tapping with a metronome occurred earlier than when tapping with music.

Moreover, there were a significant interaction of audio and MMSE, and a significant interaction of tempo stability and MMSE. Once again, these are the smallest effects (both f² < 0.001), unlikely to have any practical significance despite their being statistically significant, and they can be found in the Supplementary Information (Figures S5 and S6).

## DISCUSSION

The purpose of this study was to investigate the influence of age and NCD severity on SMS performance, quantified as tapping consistency and asynchrony. We were particularly interested in whether different subprocesses of SMS, including beat induction, beat maintenance, and error correction, might be differentially impacted by age and NCD. This differential impact could manifest in age and NCD unequally influencing SMS performance when synchronising with music versus metronomes, and with tempo-changing sequences compared to sequences with a stable tempo. We demonstrated that tapping consistency decreased with MMSE, providing evidence of the impact of neurocognitive disorders on sensorimotor abilities and thereby confirming our hypothesis. Contrary to our other hypothesis, however, the effect of MMSE did not depend on tempo stability. Additionally and as predicted, we observed a decrease in consistency with age, but only when individuals tapped with musical sequences and not with a metronome. Lastly, we found that consistency and asynchrony were differently modulated by tempo changes depending on the tempo and the type of auditory sequences. Before discussing these results in depth, it is worth noting that participants’ mean spontaneous motor tempo of 715 ms is close to what has previously been found in older adults [21,25]. More importantly, it was squarely in between the two stimulus tempi in the paced tapping task (674 ms and 741 ms). It is therefore reasonable to assume that both tempi were in the range of comfortable rates for our participants.

### Effect of NCD severity on consistency

The observed association between MMSE score and tapping consistency is in line with previous findings. A recent study by Hobeika et al [35] also found reduced consistency in people with major NCD compared to those with mild or no NCD, as well as a negative linear relationship between MMSE and consistency during an audio-visual tapping task. However, the latter result was limited to the metronome condition, whereas participants were not impaired with music. Interestingly, in our study, cognitive impairment had a global impact on consistency, affecting tapping with both metronome and music. Perhaps these differences stem from the fact that our study, which included trials with tempo changes, was more sensitive to uncovering NCD-related effects. On the other hand, the effect of NCD severity on tapping to music present here but absent in Hobeika et al.’s study might be attributed to music-induced reward. The motivating and rewarding qualities of music may boost synchronisation, resulting in more consistent tapping [65]. Perhaps Hobeika et al.’s stimuli, with original music recordings and sung lyrics, were more rewarding than our MIDI-based stimuli which did not contain lyrics and which were also repeated more often within the same experiment. Perhaps a difficulty in synchronising to music was offset by enhanced synchronicity related to reward In Hobeika et al.’s study, underscoring the relevance of selecting music for its motivating and rewarding qualities.

The finding of an NCD-related deficit in SMS at a comfortable rate is novel: Some studies have previously shown lower tapping consistency in people with NCD, but only when they had to continue tapping after an external sequence had ended [22,32] and/or when the tempo they synchronised with was far from their comfortable tempo (i.e., slower [22,34] or faster [33]. We hypothesised that the tempo-changing manipulation would be particularly difficult for more cognitively impaired people and that MMSE and tempo stability would therefore interact, but this effect was not observed in this study. It may be that the current task and its analysis pipeline, examining consistency by 15-second segments, and comparing these segments across conditions, may have been too crude, given that people only take a few taps to adapt to a new tempo [18,66,67], at least healthy participants. Additionally, it is possible that the bimodal nature of the task (audio and video) made the task easier, offsetting the difficulty introduced by the tempo changes. Finally, while MMSE was chosen as a predictor variable to capture the full spectrum of cognitive impairment, this may have resulted in reduced statistical power to detect effects, especially interaction effects, than sampling two extreme groups [68]. Additional research on rhythmic synchronisation with tempo changes is warranted, as it may provide insights into how individuals generally entrain to regularities and adapt to changes in their sensory environment.

This study, along with another recent study [35], highlights a global deficit in SMS abilities among individuals with NCD. Given the established connection between rhythmic and cognitive abilities, it can be speculated that rhythmic training may confer cognitive benefits. However, the direct transfer of benefits from musical to non-musical domains requires further investigation. There exist other neurological conditions like Parkinson’s disease, Huntington’s disease, autism spectrum disorder, attention deficit hyperactivity disorder, and dyslexia, where rhythmic deficits are prominent and rhythm-based training may offer advantages beyond the motor realm, such as on communication and executive functions [69–75]. By continuing to study SMS and its links with cognitive abilities, we may get a clearer picture of what processes may inadvertently be stimulated through rhythm-based interventions, to slow down symptoms in NCD, but also as a preventive strategy in healthy older adults [76–79]. Finally, the current results also suggest that sensorimotor problems could serve as a potential diagnostic marker of NCD, warranting inclusion in the neuropsychological evaluation process, but only as complementary tests among measures of working memory and attention, for which the link with NCD is more established.

### Interaction between audio and age on consistency

Another noteworthy result was an interaction effect of audio and age on consistency. Age negatively affected tapping consistency when people synchronised their taps with music, but not with a metronome. This observation offers a more nuanced perspective on past research that found higher consistency when tapping with metronomes compared to music [12–14] and research on the effect of age on SMS which often found null results at least with a comfortable tempo, but which rarely used music material as a stimulus, but rather metronomes (Bangert & Balota, 2012; Carment et al., 2018; Drewing et al., 2006; Duchek et al., 1994; Krampe et al., 2005; McAuley et al., 2006; Turgeon et al., 2011; Vanneste et al., 2001; but see Nagasaki et al., 1988; Thompson et al., 2015). The current findings, revealing distinct effects of age on tapping to metronomes versus tapping to music, suggest that beat maintenance and beat induction may be affected differently. Perhaps older adults experience greater impairment in beat induction processes, which are crucial for tapping with music, whereas they retain their ability in the implicit and mechanical aspects of beat maintenance, resulting in comparable performance to younger individuals when tapping with a metronome.

Previous research indicates that during movement performance in older adults, additional brain regions, specifically prefrontal areas, become active [82–84], even in situations where there are no age-related differences in performance outcomes. This suggests increased cognitive control in executing movements in older individuals. Thus, there might be a beginning decline in motor control associated with aging, which people compensate for by employing extra neural and cognitive resources, leading them to achieve performance levels comparable to those of younger individuals when the task is simple, such as metronome tapping in this study. However, in tasks that demand higher-level representations and/or executive control such as bimanual [85–88] and sequential [89] tapping, or having to rapidly extract the beat from a musical sequence such as in this study, these compensatory mechanisms might not be sufficient, leading to age-related differences in performance in these more complex tasks. The global effect of MMSE on consistency discussed in the previous section may also imply that people with NCD do not engage in compensatory mechanisms as efficiently as healthy older adults, or that this compensation is not sufficient to mask differences in performance even on simpler tasks like tapping with a metronome. For future research, it is crucial to use stimuli with varying levels of complexity, as in this study, to discern the factors that yield observable performance differences.

### Interactions between audio, tempo, and tempo stability on consistency, and on asynchrony

With regards to consistency, a three-way interaction emerged. Besides the expected decrease in consistency following tempo changes and the lower consistency observed with music compared to metronomes, we noted that consistency was affected by tempo only in one specific circumstance. Specifically, if the auditory stimulus was a metronome with a shifting tempo, consistency was higher when the tempo was slow (i.e., following a deceleration) than when it was fast (i.e., following an acceleration). This complements existing literature which has reported decelerations as being easier to detect [18] and more easily adapted to [90–93].

Regarding asynchrony, we replicated one of the most stable findings in the SMS literature, the mean negative asynchrony [5,8,12,13,16,50,94]: Regarding the trials in which the tempo was stable, patients’ taps preceded the beats by several tens of milliseconds when synchronising with a metronome, whereas they tended to tap close to the beat when tapping with music. In trials following a tempo change, we would expect taps to be late after an acceleration and anticipated after a deceleration, relatively to the value of asynchrony in the corresponding stable condition, at least for a few taps [95,96]. Our results confirm this response pattern in the musical trials, and after the deceleration of a metronomic tempo. However, after an acceleration with a metronome, asynchrony remains negative and indeed close to the asynchrony during a fast stable tempo. This pattern may indicate that people adapt more quickly or efficiently to accelerating than decelerating metronomes, which would be in contrast to the result we observe for consistency. However, it may also be that a deceleration is indeed more easily perceived, as in past studies, but accompanied by an overcorrection response, leading to the large observed negative asynchrony. Finally, since only one piece of music was used in this study, the observed asymmetry may also arise from idiosyncrasies specific to this particular piece, especially considering that the original song contains considerable tempo changes that may have influenced participants’ expectations of the temporal structure.

### Implications for music-based interventions

The results highlight that motor and cognitive skills may be tightly linked, indicating the potential of rhythm-based interventions to stimulate non-motor domains, such as working memory, executive functions, language, and socio-emotional functioning, presenting a promising avenue for improving the quality of life in individuals with NCDs. The current findings are relevant to how interventions may be tailored to a person’s cognitive status. Considering that individuals with lower cognitive functioning may have difficulties in synchronising movements to auditory stimuli, particularly those that are not intrinsically motivating or rewarding, it is essential to adapt music-based interventions based on cognitive ability and carefully select appropriate stimuli. One may consider using stable and predictable beats, potentially including metronomes or music with high beat clarity when working with older adults, given the age-related decline in beat induction demonstrated here. The observed reduction in consistency when introducing tempo changes could serve as an argument for adaptive programmes, starting with simpler, stable tempi and gradually introducing more complex rhythms to ensure task engagement and build rhythmic skills progressively. While our research and its implications for rhythm-based interventions are focused on simple, unimanual tapping, it is essential to note that music-based interventions requiring finer motor control may specifically engender cognitive benefits [97]. While this study did not compare audio-visual stimuli with purely auditory stimuli, the high levels of performance observed here suggest that visual cues of any kind may enhance synchronisation. Finally, non-musical cognitive training could be intertwined with musical exercises, mutually enhancing each other’s effectiveness.

### Limitations

Our sample included individuals with NCD of diverse origins, predominantly AD, vascular NCD, and NCD of mixed aetiology. While this sample is likely representative of the general population of individuals with NCD, the limited numbers within each subgroup did not allow us to explore differences between various aetiologies, which presents an interesting avenue for future research. In fact, we are aware of only one study [34] that compared sensorimotor synchronisation abilities across different NCD groups and identified differences between AD and frontotemporal NCD.

We recognised the importance of good hearing and vision for our experiment, screening out potential participants with impairments or those who did not have the necessary aids with them. However, we did not conduct formal audiometry or visual acuity tests, leaving the possibility that performance variations could be attributed in part to differences in hearing and visual abilities, considering the common prevalence of hearing loss [98] and visual impairment [99,100] in older adults.

It is worth repeating that we deliberately chose to use audio-visual stimuli to synchronise to, a manipulation deemed necessary to maintain participants’ engagement and motivation throughout the task. Nevertheless, this prevented us from assessing the degree to which participants relied on auditory versus visual information, or how performance might be affected if individuals had access to information from only one modality, which presents an interesting direction for future investigation.

### Concluding remarks

This study highlights two primary findings. The first is an influence of MMSE score on tapping consistency, irrespective of audio stimulus type and of the presence or absence of tempo changes, suggesting an effect of NCD severity on the ability to maintain a steady rhythm. Two possible mechanisms could explain this, which are not mutually exclusive. Firstly, neural reorganising over the course of the disorder may increasingly engage non-motor areas to sustain performance during a simple motor task, indicating a shift towards more cognitive and effortful processing of rhythm. Secondly, even simple metronome tapping may require some degree of attention and working memory, albeit less than tapping with music. Healthy ageing individuals may therefore maintain a consistent level of performance when tapping with a metronome, whereas in more cognitively impaired individuals with a lower MMSE, the impairment of attention and working memory is severe enough to significantly hinder performance, even in tapping with a simple metronome, arguably the simplest form of SMS. However, it is important to acknowledge the importance of tempo changes in half of the trials. While the statistical analysis indicates that the impact of cognitive impairment held for both conditions (with and without tempo changes), it may still be that the primary difficulty in this study might have arisen from the presence of tempo changes in half of the trials, even though the difference in decline of consistency as a function of MMSE across the two levels of tempo stability was not large enough to yield statistical significance. Although we did not observe an interaction between tempo stability and MMSE in this study, the current results do not eliminate the possibility that individuals with NCD might experience specific difficulties in adapting to tempo changes. The involvement of working memory, attention, and cognitive flexibility in error correction could still play a role, warranting further investigation. The second result shows an age-related decline in consistency during SMS but only when tapping with music, whereas consistency remains stable when tapping with a metronome. This observation implies that beat induction, a process especially relevant for perceiving the underlying beat in musical sequences, is affected in healthy ageing, potentially indicating a beginning decline of attention and working memory. Beat maintenance, on the other hand, may be relatively spared.

In conclusion, this research emphasises the importance of sensorimotor impairment as a symptom in NCD. The findings suggest that motor and cognitive skills may be tightly linked, implying that deficits in one domain may potentially impact the other. This underplay underlines the potential for rhythm-based training to inadvertently stimulate non-motor domains, such as working memory, executive functions, language, and socio-emotional functioning, presenting a promising avenue for enhancing the quality of life in individuals living with NCDs. These insights provide a foundation for continued research and therapeutic interventions aimed at enhancing well-being in healthy and pathological ageing by targeting the sensorimotor domain. The possibility for therapeutic approaches are vast, ranging from group drumming [101] to remote interventions using mobile devices [102].

## ACKNOWLEDGEMENTS

This project has received funding from the European Union’s Horizon 2020 research and innovation programme under the Marie Skłodowska-Curie grant agreement No 847568. Moreover, this study was supported by the French government through the Programme Investissement d’Avenir (I-SITE ULNE / ANR-16-IDEX-0004 ULNE) managed by the Agence Nationale de la Recherche.

The authors thank Ivan Schepers of Ghent University for help in the material’s development, and the musician Sotirios Sideris with whom the stimuli were developed. We furthermore thank the geriatrician Jean Roche, the psychologists Anita Clercx, Sylvie Schoenenburg, and Laurence Grymonprez, and the entire dedicated staff at the day hospital Les Bateliers in Lille. Finally, we thank all participants involved in this study.

During the preparation of this work the authors used the large language model ChatGPT [103] in order to improve the flow and readability of the writing. After using this tool, the authors reviewed and edited the content as needed and take full responsibility for the content of the publication.

## DATA AVAILABILITY STATEMENT

The data supporting the findings and a script to analyse them are openly available at https://osf.io/78k46/?view_only=9e15fa4ac33d49e1aff47bd609c305ab.

