## Supplementary Material for "Sensorimotor Impairment in Ageing and Neurocognitive Disorders: Beat Synchronisation and Adaptation to Tempo Changes"

SUPPLEMENTARY TABLES AND FIGURES

**Table S1**

*Musician's performance. When recording the videos of the musician tapping synchronously with the auditory sequences, inter-tap interval and coefficient of variation were verified to ensure that the musician tapped with minimal error.*

| | Inter-tap interval (ms) | Coefficient of variation |
|---|---|---|
| 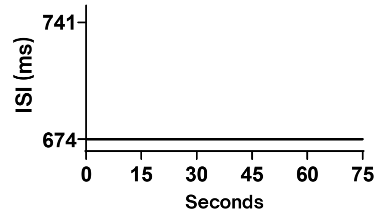  | 8.90 | 0.40 |
| 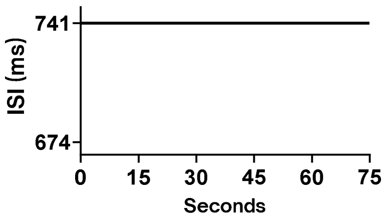  | 1.01 | 0.05 |
| 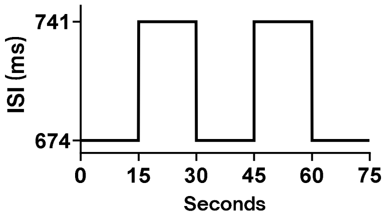  | -0.11 | 0.00 |
| 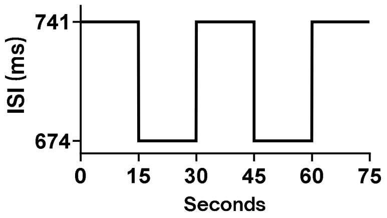  | -0.85 | 0.02 |

**Table S2**

*Demographic and clinical information of patients who withdrew from the study during the experiment.*

| | N | Median [first quartile, third quartile] or frequencies (%) | Range |
|---|---|---|---|
| Age | 7 | 88 [84, 94] | 81-95 |
| Gender (women) | 7 | 5 (71%) | |
| Diagnosis | 6 | | |
| Major NCD | | 6 (100%) | |
| Minor NCD | | 0 (0%) | |
| No NCD | | 0 (0%) | |
| Years of education | 6 | 9 [7, 14] | 5-14 |
| Musical expertise (out of 28) | 7 | 3 [2, 4] | 0-4 |
| MMSE (out of 30) | 7 | 21 [20, 25] | 17-29 |
| ADL (out of 6) | 7 | 5.5 [4, 6] | 2-6 |
| IADL (out of 4) | 7 | 1 [0, 2] | 0-4 |
| GDS-SF (out of 15) | 6 | 4 [3, 7] | 2-7 |
| GAI-SF (out of 5) | 6 | 1 [1, 1] | 0-4 |

NCD = neurocognitive disorder; MMSE = Mini-Mental State Examination; ADL = activities of daily living; IADL = instrumental activities of daily living; GDS-SF = Geriatric Depression Scale - Short Form; GAI-SF = Geriatric Anxiety Inventory - Short Form

**Table S3**

*Model summary and significance tests: consistency*

| | Wald $\chi^2$ | df | p | $f^2$ |
|---|---|---|---|---|
| Sex | 0.236 | 1 | .627 | 0.006 |
| Years of education | 0.446 | 1 | .504 | 0.022 |
| Musical expertise | 1.309 | 1 | .253 | 0.035 |
| Condition order | 1.502 | 1 | .220 | 0.002 |
| Audio | 86.814 | 1 | < .001* | 0.261 |
| Tempo | 0.210 | 1 | .647 | < 0.001 |
| Tempo stability | 794.225 | 1 | <.001* | 0.217 |
| Segment | 3.652 | 1 | .056 | < 0.001 |
| Age | 1.814 | 1 | .178 | 0.010 |
| MMSE | 4.057 | 1 | .044* | 0.032 |
| Audio:Tempo | 1.100 | 1 | .294 | < 0.001 |
| Audio:Tempo stability | 2.525 | 1 | .112 | < 0.001 |
| Audio:Segment | 11.867 | 1 | .001* | 0.005 |
| Tempo:Tempo stability | 0.162 | 1 | .688 | < 0.001 |
| Tempo:Segment | 6.649 | 1 | .010* | 0.003 |
| Tempo stability:Segment | 5.961 | 1 | .015* | 0.003 |
| Audio:Age | 7.056 | 1 | .008* | 0.003 |
| Tempo:Age | 0.238 | 1 | .626 | < 0.001 |
| Tempo stability:Age | 8.829 | 1 | .003* | 0.001 |
| Audio:MMSE | 0.104 | 1 | .747 | < 0.001 |
| Tempo:MMSE | 0.017 | 1 | .896 | < 0.001 |
| Tempo stability:MMSE | 1.085 | 1 | .298 | < 0.001 |

| | | | | |
|---|---|---|---|---|
| Audio:Tempo:Tempo stability | 5.702 | 1 | .017* | 0.003 |
| Audio:Tempo:Segment | 1.394 | 1 | .238 | < 0.001 |
| Audio:Tempo stability:Segment | 1.351 | 1 | .245 | < 0.001 |
| Tempo:Tempo stability:Segment | 0.003 | 1 | .957 | < 0.001 |
| Audio:Tempo:Age | 0.774 | 1 | .379 | < 0.001 |
| Audio:Tempo stability:Age | 1.262 | 1 | .261 | < 0.001 |
| Tempo:Tempo stability:Age | 0.423 | 1 | .515 | < 0.001 |
| Audio:Tempo:MMSE | 4.082 | 1 | .043* | 0.002 |
| Audio:Tempo stability:MMSE | 0.008 | 1 | .928 | < 0.001 |
| Tempo:Tempo stability:MMSE | 0.451 | 1 | .502 | < 0.001 |
| Audio:Tempo:Tempo stability:Segment | 0.627 | 1 | .429 | < 0.001 |
| Audio:Tempo:Tempo stability:Age | 1.542 | 1 | .214 | 0.001 |
| Audio:Tempo:Tempo stability:MMSE | 3.767 | 1 | .052 | < 0.001 |

Significant results are highlighted in **bold**.

*Abbreviation*: MMSE = mini-mental state examination

**Table S4**

*Model summary and significance tests: asynchrony*

| | Wald $\chi^2$ | df | p | $f^2$ |
|---|---|---|---|---|
| Sex | 1.542 | 1 | .214 | 0.004 |
| Years of education | 0.224 | 1 | .636 | < 0.001 |
| Musical expertise | 1.546 | 1 | .214 | 0.004 |
| Condition order | 0.277 | 1 | .599 | < 0.001 |
| Audio | 454.145 | 1 | < .001* | 0.112 |
| Tempo | 253.003 | 1 | < .001* | 0.136 |
| Tempo stability | 4.090 | 1 | .043* | 0.012 |
| Segment | 90.031 | 1 | < .001* | 0.018 |
| Age | 1.912 | 1 | .167 | 0.004 |
| MMSE | 2.912 | 1 | .088 | 0.001 |
| Audio:Tempo | 77.939 | 1 | < .001* | 0.033 |
| Audio:Tempo stability | 0.006 | 1 | .938 | 0.013 |
| Audio:Segment | 9.327 | 1 | .002* | 0.003 |
| Tempo:Tempo stability | 246.478 | 1 | < .001* | 0.139 |
| Tempo:Segment | 101.387 | 1 | < .001* | 0.068 |
| Tempo stability:Segment | 12.899 | 1 | < .001* | 0.004 |
| Audio:Age | 0.471 | 1 | .492 | 0.001 |
| Tempo:Age | 2.008 | 1 | .156 | < 0.001 |
| Tempo stability:Age | 3.241 | 1 | .072 | < 0.001 |
| Audio:MMSE | 4.614 | 1 | .032* | 0.001 |
| Tempo:MMSE | 1.062 | 1 | .303 | < 0.001 |
| Tempo stability:MMSE | 4.374 | 1 | .036* | < 0.001 |

| | | | | |
|---|---|---|---|---|
| Audio:Tempo:Tempo stability | 15.151 | 1 | < .001* | 0.015 |
| Audio:Tempo:Segment | 8.946 | 1 | .003* | 0.003 |
| Audio:Tempo stability:Segment | 35.843 | 1 | < .001* | 0.008 |
| Tempo:Tempo stability:Segment | 49.626 | 1 | < .001* | 0.055 |
| Audio:Tempo:Age | 1.434 | 1 | .231 | < 0.001 |
| Audio:Tempo stability:Age | 3.035 | 1 | .081 | 0.001 |
| Tempo:Tempo stability:Age | 3.521 | 1 | .061 | 0.001 |
| Audio:Tempo:MMSE | 0.056 | 1 | .813 | < 0.001 |
| Audio:Tempo stability:MMSE | 0.284 | 1 | .594 | < 0.001 |
| Tempo:Tempo stability:MMSE | 0.549 | 1 | .459 | 0.002 |
| Audio:Tempo:Tempo stability:Segment | 27.592 | 1 | < .001* | 0.015 |
| Audio:Tempo:Tempo stability:Age | 2.467 | 1 | .116 | 0.001 |
| Audio:Tempo:Tempo stability:MMSE | 1.516 | 1 | .218 | 0.001 |

Significant results are highlighted in **bold**.

*Abbreviation*: MMSE = mini-mental state examination

**Figure S1**

*Within-subject variables audio, tempo, and tempo stability, emerging from the eight experimental trials*

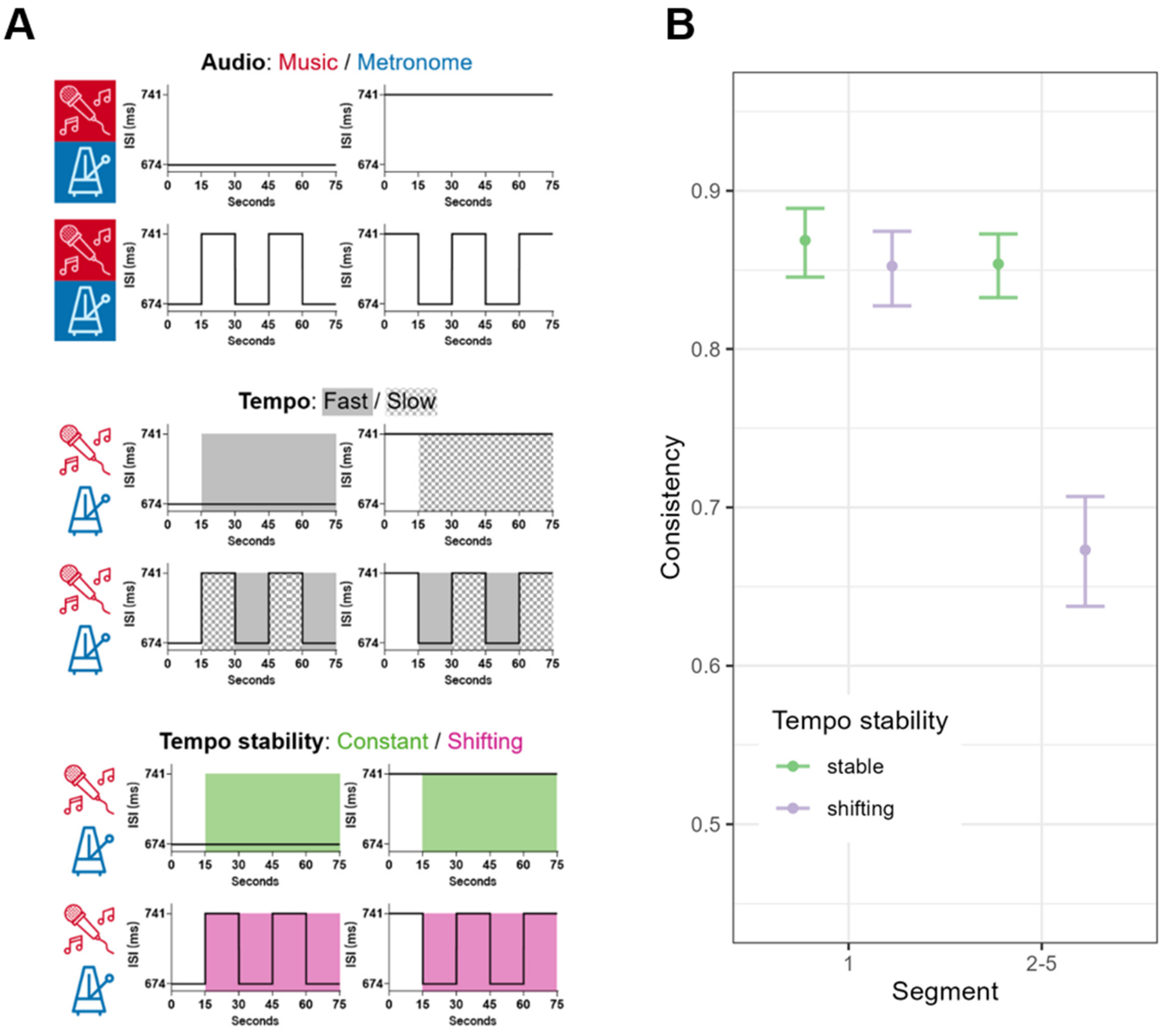


*Note.* (A) Analysing tapping consistency and asynchrony over 15-second segments allowed us to independently study the effects of audio (music vs. metronome), tempo (fast vs. slow), and tempo stability (constant vs. shifting). (B) To provide an additional verification, consistency was calculated individually for the first segment of each trial and for segments 2-5 of each trial, highlighting the drop in performance as a result of changes in tempo.

**Figure S2**

*Distribution of MMSE*

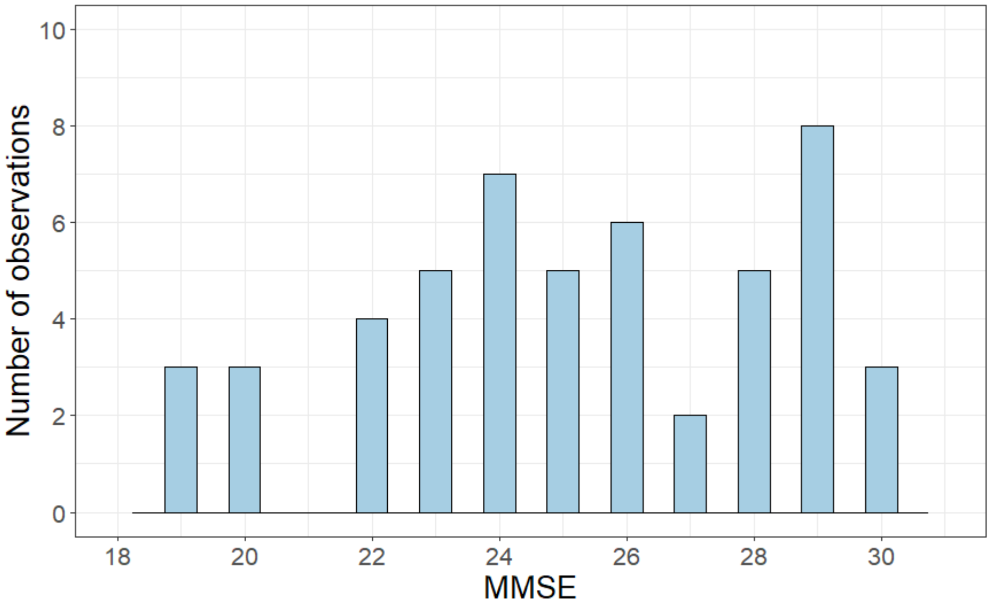


*Note.* MMSE, Mini-Mental State Examination

**Figure S3**

*Effects of age and tempo stability and consistency*

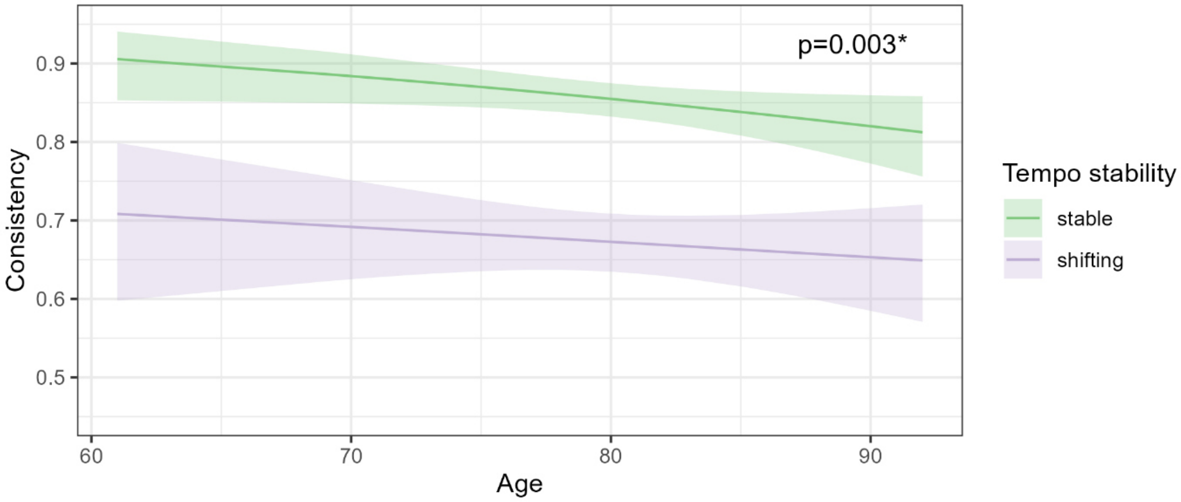

**Figure S4**

*Effects of audio, tempo, and MMSE on consistency (p < .001)*

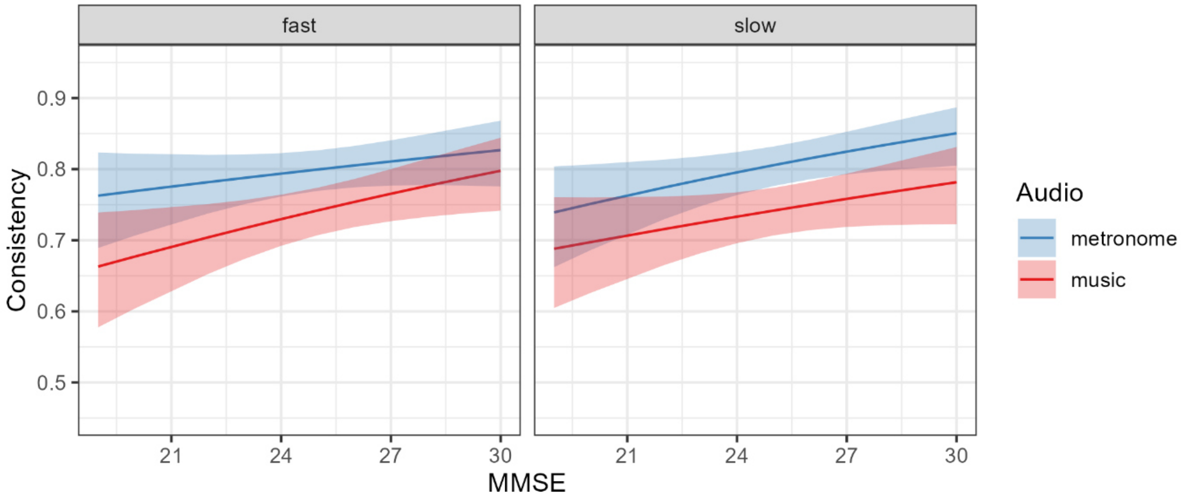


**Figure S5**

*Effects of audio and MMSE on asynchrony*

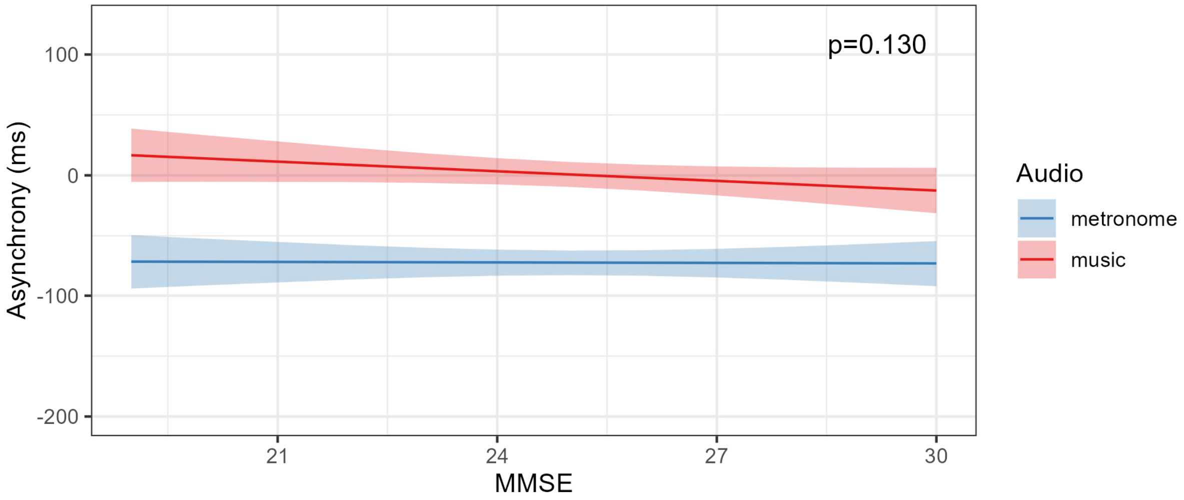


*Note*. This graph was created by taking estimated marginal means and standard deviations of a linear mixed model with non-transformed data (whereas the statistical tests were done with a model in which the response variable, asynchrony, was transformed by taking the cubic root of its absolute value and multiplying it with its original sign).

**Figure S6**

*Effects of tempo stability and MMSE on asynchrony*

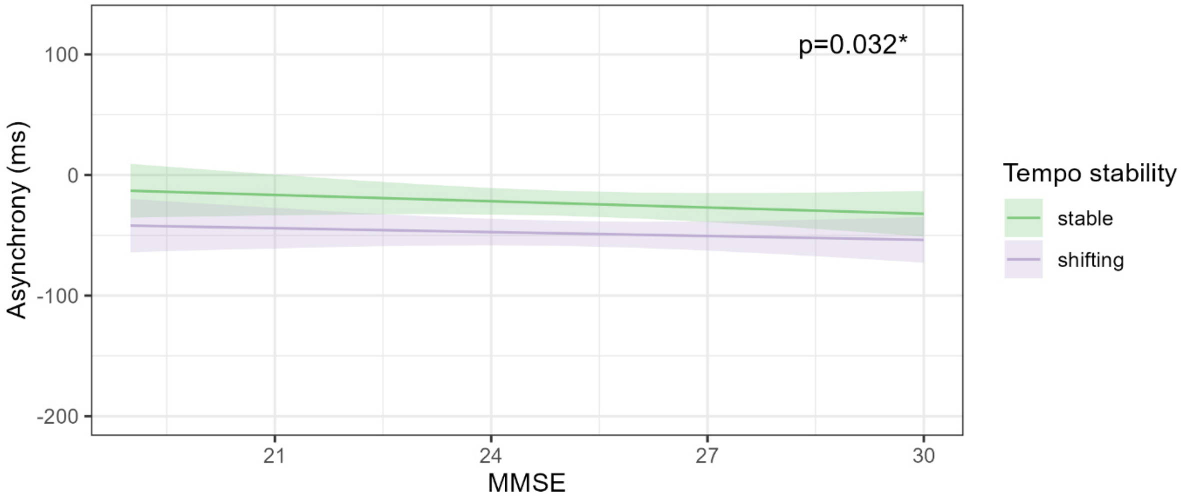


*Note*. This graph was created by taking estimated marginal means and standard deviations of a linear mixed model with non-transformed data (whereas the statistical tests were done with a model in which the response variable, asynchrony, was transformed by taking the cubic root of its absolute value and multiplying it with its original sign).
